# Feeling the music: Audiotactile encoding of temporal structure in the human brain

**DOI:** 10.64898/2026.01.17.700057

**Authors:** Giulio Degano, Ambra Ferrari, Uta Noppeney

**Author notes:** Corresponding author (AF). These authors contributed equally to this work and share first authorship (Giulio Degano, Ambra Ferrari).

## Abstract

In everyday situations, like a rock party or an organ concert, we feel music vibrating through our bodies. How do these vibrotactile signals influence music processing? How do they aid auditory scene analysis? Combining psychophysics, fMRI and time-resolved EEG decoding, this work reveals how the brain encodes the temporal structure of music (beat and envelope) across audition and touch and uses this information to guide multisensory integration and segregation in simple and more complex perceptual scenes. Participants experienced monophonic and polyphonic piano pieces through auditory, vibrotactile and audiotactile stimulation. Vibrotactile signals improved the detection of a brief target embedded in music, establishing the functional relevance of audiotactile integration in naturalistic settings. fMRI and EEG multivariate decoding revealed that auditory and tactile beat information converged in planum temporale and parietal operculum, albeit through distinct neural dynamics and representations. Superior temporal cortices reliably encoded envelope information from audition, but only weakly from touch. Nevertheless, vibrotactile signals significantly enhanced neural encoding of auditory beat as early as 100 ms, and envelope representations from 250 ms onward. These encoding benefits were associated with superadditive interactions in primary auditory cortex, where tactile signals sharpen and amplify auditory envelope representations. In complex polyphonic music, touch further amplified the segregation and encoding of temporally coherent auditory streams. Our findings highlight the important, yet largely unexplored influence of touch on auditory processing, enriching music perception and supporting auditory scene analysis in real-world environments.

## Introduction

Imagine attending an organ concert or dancing at a rock festival. Not only do you enjoy listening to your favorite music, but you also feel the vibrations produced by the music resonating through your body. Audition and touch have traditionally been regarded as complementary senses: audition as a “far sense” allows us to perceive sources in the distance or outside our field of view, while touch as a “near sense” typically requires direct physical contact with objects. However, touch can also extend its reach by sensing mechanical pressure waves that travel through the air (Sathian & Lacey, 2022; Soto-Faraco & Deco, 2009). Most notably, loud music produces mechanical pressure waves that can be heard as they impinge on the eardrum and felt as they stimulate vibrotactile sensors throughout the entire body.

Despite a growing body of research on multisensory integration, little is known about how the brain merges information from audition and touch. Audiotactile interactions have been observed in non-human primates in the caudal belt, a region in posterior secondary auditory cortices (Kayser et al., 2005, 2008; Schroeder et al., 2008), and in humans even in primary auditory cortices (Hoefer et al., 2013; Schürmann et al., 2006). Neurophysiological work revealed a temporal cascade of audiotactile interactions arising in evoked response potentials (Foxe et al., 2000; Molholm et al., 2002; Zumer et al., 2020), inter-trial coherence (Zumer et al., 2020) and induced oscillations (Lakatos et al., 2007; Leonardelli et al., 2015; Zumer et al., 2020). However, these studies used artificial, simple transient stimuli that are far removed from everyday experience (Nastase et al., 2020; Sonkusare et al., 2019). Music perception offers a unique opportunity to study how the brain combines auditory and tactile signals in naturalistic dynamic scenes (Bizley et al., 2016; Campbell, 2007; Shamma et al., 2011).

Music includes several complex time-varying features, such as beat and envelope, which evolve with distinct temporal structures. While beat defines the tempo as a steady regular pulse, envelope represents changes in sound amplitude across time. Previous research on auditory music perception has shown that both beat and envelope processing primarily activate the posterior superior temporal gyrus and planum temporale (BA 42/22) with additional activations observed in premotor cortex and posterior cerebellum (Di Liberto et al., 2020; Giraud et al., 2000; Grahn et al., 2011; Grahn & Brett, 2007; Large et al., 2015; Vuust et al., 2022). Importantly, although audition is the primary sense for music perception, touch can also provide information about both the music’s beat and envelope, thereby increasing the immersive nature of music experience (Hove et al., 2020; Huang et al., 2012; Tranchant et al., 2017). From the computation perspective of Bayesian Causal Inference, the brain should combine signals that arise from common causes, but segregate those from separate causes (Noppeney, 2021; Shams & Beierholm, 2022). Temporal correspondences, including both beat and envelope features, provide critical cues informing the brain whether touch and auditory signals come from common causes and should thus be integrated. This raises the critical question of how the brain extracts and combines beat and envelope information from audition and touch to guide multisensory integration and segregation in simple and more complex perceptual scenes (Bizley et al., 2016; Campbell, 2007; Shamma et al., 2011).

We addressed this outstanding question in a multi-day study combining psychophysics, functional magnetic resonance imaging (fMRI) and electroencephalography (EEG). To mimic the music scenarios described above, we modulated the amplitude of the vibrotactile signals to match the beat and acoustic envelope of one of the two possible voices of polyphonic counterpoint piano pieces, custom-composed by a professional musician. Observers were presented with one (i.e. monophonic) or two different (i.e. polyphonic) voices of the piano pieces in auditory, tactile or audiotactile contexts. While all music pieces shared the same beat, we manipulated the congruency of envelope information in the audiotactile conditions. Congruent or incongruent envelope-modulated vibrotactile signals were applied to participants’ index fingertips. In the psychophysics experiment, we evaluated whether a concurrent tactile stream boosted the participants’ ability to detect a target event embedded within an auditory monophonic or polyphonic stream. During the fMRI and EEG recordings, participants were engaged in a visual detection task to monitor their attention while avoiding confounding influences of task or overt responses on audiotactile processing.

This study allowed us to address three key questions. First, does the brain encode beat and envelope information from purely auditory and vibrotactile signals via shared or distinct neural representations, and how do these evolve across time? Second, does a concurrent vibrotactile signal influence the encoding of auditory beat and envelope information, and how are these processes modulated by whether auditory and tactile signals share temporal envelope information? Finally, does a tactile signal enhance the segregation and encoding of the temporally correlated auditory envelope in more complex polyphonic scenarios? Using psychophysics, we first examined whether touch produces behaviorally relevant effects. Importantly, we then investigated how these effects arise in terms of cortical architectures and temporal dynamics through the combination of whole-brain fMRI and time-resolved EEG decoding.

Using psychophysics, we demonstrate that a tactile stream carrying congruent beat and envelope information boosts the ability to detect a target event embedded within a monophonic or polyphonic auditory stream. Combining analyses of regional BOLD-responses with time-resolved EEG decoding, we reveal that touch clearly conveys beat information but is rather poor in contributing envelope information. Yet, both tactile beat and envelope information modulate auditory processing and support auditory scene analysis in bilateral temporo-opercular and posterior insular regions. Collectively, our findings suggest that multisensory interactions across the auditory cortical hierarchy support the automatic tracking of complex streams of information in our natural world.

## Results

The study included a psychophysics experiment followed by an fMRI and EEG experiment on separate days. During the experiments, we presented counterpoint piano pieces custom-made by a professional composer. For each music piece, envelope and frequency features were extracted from the audio signal and used to synthesize the tactile signal (Fig 1A). Congruent and incongruent audiotactile stimuli respectively combined corresponding or distinct envelope information. We ensured that incongruent stimuli minimized the temporal correlations between auditory and tactile signals (Fig 1B). To verify that participants were able to discriminate between audiotactile congruent and incongruent stimuli reliably, all experiments set an accuracy threshold on their congruency judgments as an a priori inclusion criterion (Fig 1C; see section “Inclusion criterion: congruency judgement”). In the psychophysics experiment, two volunteers were excluded from the study due to poor accuracy in distinguishing between audiotactile congruent and incongruent music streams. The included participants showed highly accurate responses (d′ = 3.27 ± 0.05; *bias* = -0.01 ± 0.04, across-participants’ mean ± SEM). In the fMRI and EEG experiments, all participants performed accurately (d′ = 3.16 ± 0.10; *bias* = 0.01 ± 0.04, across-participants’ mean ± SEM) and were therefore included in the analyses and results.

**Figure 1.**
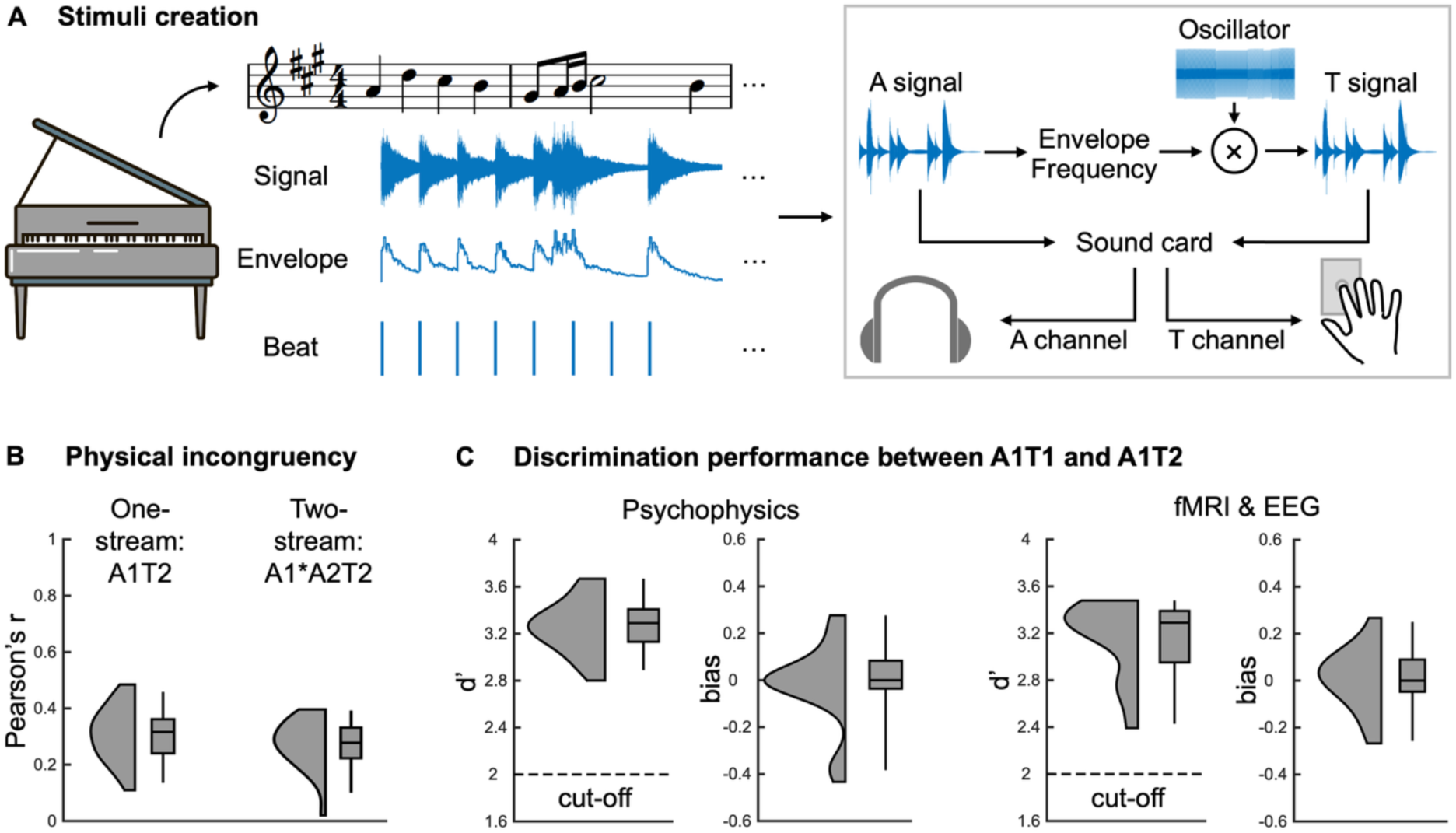
Stimuli validation. **A)** Auditory stimuli were counterpoint piano pieces custom-made in collaboration with a professional composer. Music features (envelope and frequency) were extracted from the audio signal and used to synthesize the tactile signal through a sinusoidal oscillator. We thereby obtained corresponding streams in audition and touch for each music piece. **B)** Probability density graphs and boxplots of the across streams’ Pearson’s correlation coefficient (r) between audio and tactile streams in the one-stream (A1T2) and two-stream (A1*A2T2) incongruent conditions. Incongruent conditions were created by minimizing the envelope correlation between streams via 10,000 paired permutations and selecting pairs that minimized the correlation score. **C)** Participants were included in the study based on their ability to discriminate between audiotactile congruent (A1T1) and incongruent (A1T2) streams. Probability density graphs and boxplots of the across participants’ d’ and bias show highly accurate congruency judgements for all included participants.

In the psychophysics experiment, participants were exposed to one (i.e. monophonic) or two (i.e. polyphonic) voices of the piano pieces in unisensory auditory or audiotactile contexts (Fig 2A). In a yes-no target detection task, they reported whether a brief 2Hz sinusoidal modulation of envelope called “tremolo” was present in the auditory and/or tactile streams via pedal press. The experiment comprised six conditions: one-stream auditory (A1); one-stream tactile (T1); one-stream audiotactile congruent (A1T1); two-stream auditory (A1A2); two-stream audiotactile with target in the congruent streams (A1*A2T1) or in the incongruent stream alone (A1A2*T1).

**Figure 2.**
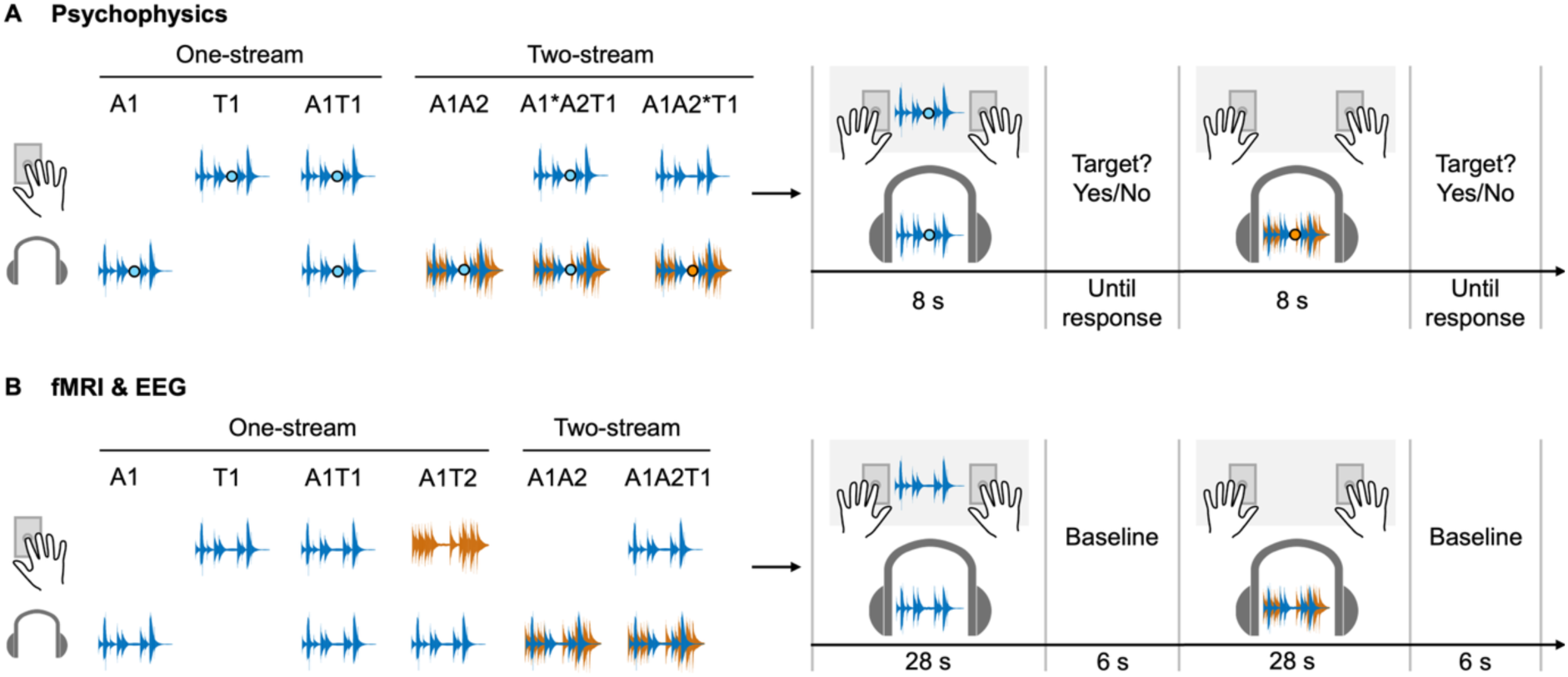
Design and procedure. **A)** In a psychophysics experiment, participants were exposed to one (i.e. monophonic) or two (i.e. polyphonic) voices of the piano pieces in auditory, tactile or audiotactile contexts. In a yes-no target detection task, they reported whether a target (“tremolo”, here represented by a dot) was present in the music stream. The experiment comprised six conditions: one-stream auditory (A1); one-stream tactile (T1); one-stream audiotactile congruent (A1T1); two-stream auditory (A1A2); two-stream audiotactile with target in the congruent streams (A1*A2T1) or in the incongruent stream (A1A2*T1). After stimulus presentation (8 s), participants reported via pedal press whether a target was present or not. Each new trial started after the participant’s response. **B)** The fMRI and EEG experiments used an identical blocked design with the same six conditions. Each condition was presented separately in a stimulation block (duration: 28 s) followed by a baseline block (duration: 6 s in fMRI, 2 s in EEG). Observers were engaged in a visual oddball task: they reported via pedal press the occasional appearance of full-screen light-grey flashes (duration: 50 ms) interspersed during stimulation blocks (none to 2 flashes per block).

The fMRI and EEG experiments included six experimental conditions (Fig 2B): one-stream auditory (A1), one-stream tactile (T1), one-stream audiotactile congruent (A1T1), one-stream audiotactile incongruent (A1T2), two-stream auditory (A1A2) and two-stream audiotactile (A1A2T1). In the two-stream audiotactile (A1A2T1) condition, the tactile envelope was congruent to one of the two auditory streams in the polyphonic piece (e.g. T congruent to A1 is denoted as A1*A2T1). To maintain attention and avoid response- and task-related confounds, participants reported the rare occurrences of brief full-screen light-grey flashes (i.e. in a third sensory modality) interspersed during the stimulation blocks (none to 2 flashes per block) via pedal press. Participants’ high detection performance confirmed their attentiveness throughout the fMRI and EEG experiments (EEG: 89.32% ± 3.21, fMRI: 84.03% ± 4.21, across-participants’ mean ± SEM % hits).

### Encoding of beat and envelope from audition and touch

First, we asked to what extent and how the brain encodes beat and envelope information extracted from purely auditory (A1) and vibrotactile (T1) signals. Using fMRI, we assessed the topographical convergence of beat and envelope representations using the “conjunction null” conjunction analysis [A1 > baseline] ∩ [T1 > baseline]. We identified common activations for auditory and tactile beat processing in the bilateral parietal opercula and plana temporalia; conversely, we observed significant activations for envelope processing only for auditory but not for touch stimulation. As a consequence, there were no significant activations for envelope processing shared across audition and touch (Fig 3A; Table 1). However, please note that we found limited activations in bilateral parietal opercula and plana temporalia also for envelope processing, shared across audition and touch, when applying a much lower significance threshold (p < 0.05 uncorrected peak-level).

**Figure 3.**
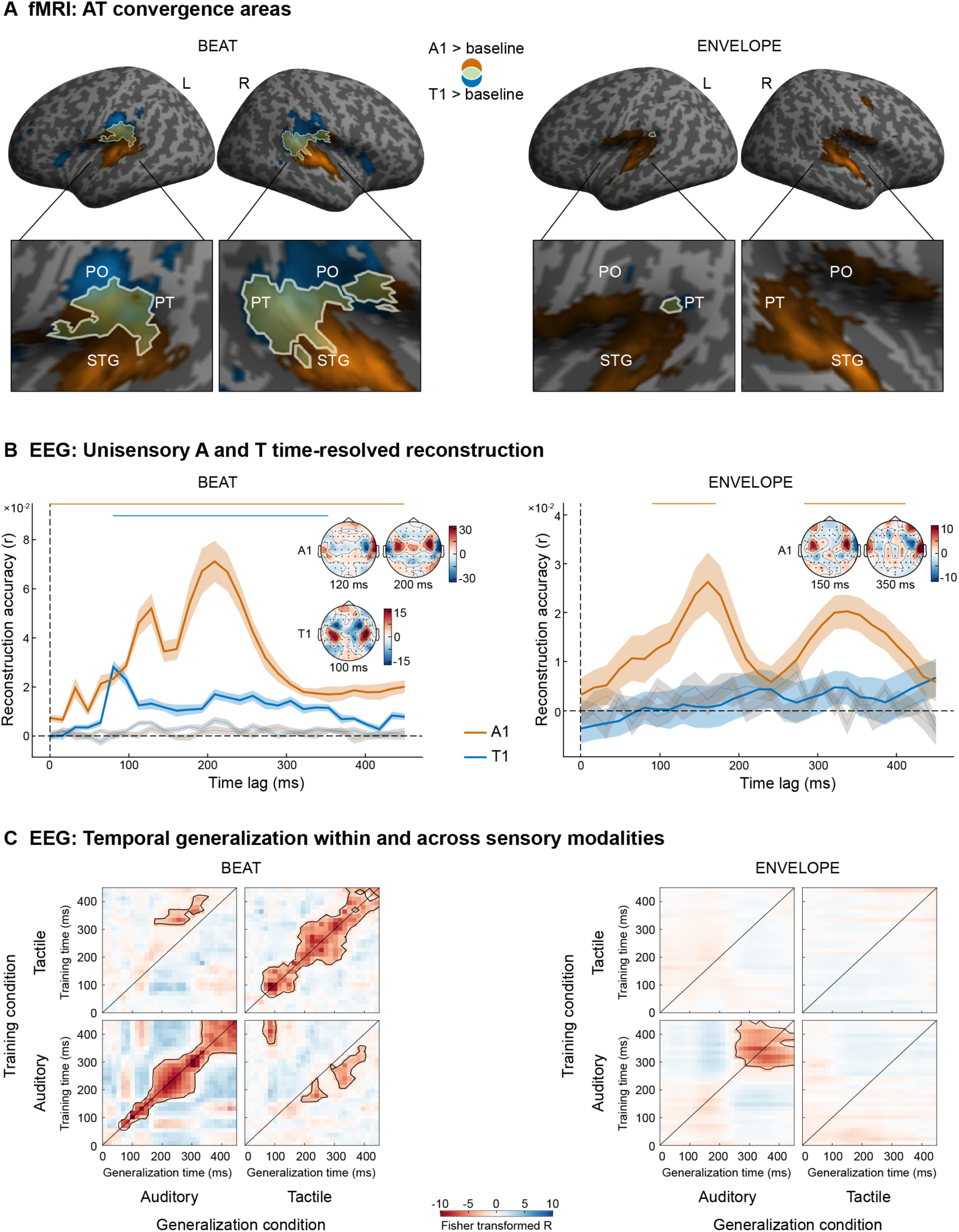
Encoding of music temporal structure from audition and touch. **A)** Audiotactile convergence effect for beat (left) and envelope (right). Increased activations for [A1 > baseline] in orange, [T1 > baseline] in blue and their overlap in yellow, rendered on an inflated canonical brain (p < 0.001 uncorrected at peak level for visualization purposes, extent threshold k > 0 voxels). White contours represent the formal “conjunction null” conjunction [A1 > baseline] ∩ [T1 > baseline]. PO, parietal operculum; PT, planum temporale; STG, superior temporal gyrus. A1, one-stream auditory; T1, one-stream tactile. **B)** Across-participants’ average time-resolved reconstruction of beat (left) and envelope (right) features for one-stream auditory (A1, orange) and one-stream tactile (T1, blue), compared with baseline (feature-shuffled) results (grey). Plots show mean (line) ± SEM (shaded area). Reconstruction accuracy is quantified in terms of Fisher-z-transformed Pearson’s correlation (r) between the reconstructed and true feature time-series. Horizontal lines show statistically significant temporal clusters (p < 0.05 corrected for multiple comparisons). **C)** Temporal generalization matrices showing reconstruction accuracy within and across sensory, i.e. auditory and tactile, modalities for beat (left) and envelope (right), by factorially manipulating the decoding scheme in a 2 (auditory versus tactile) × 2 (training versus testing) design. Each matrix shows the decoding accuracy for each training (y-axis) and testing (x-axis) time point. To assess whether auditory and tactile information rely on shared or distinct representations, we trained separate decoding models for auditory and tactile modalities and applied them to the same or opposite sensory modalities. Decoding accuracy is quantified by Pearson’s correlation between the true and the decoded information (beat or envelope, respectively). The diagonal lines indicate where the training and testing times are equal (i.e., the time-resolved decoding accuracies). The black lines encircle clusters with decoding accuracies that were significantly better than chance at p < 0.05 corrected for multiple comparisons.

**Table 1.** fMRI results: Audiotactile convergence. p-values are FWE-corrected at the cluster level for multiple comparisons within the entire brain, with an auxiliary uncorrected peak-level threshold of p < 0.001. A: auditory; T: tactile; L: left; R: right.

| Brain regions | MNI coordinates<br>(mm) |  |  | Cluster size<br>(voxels) | z-score<br>(peak) | p FWE-corrected<br>(cluster) |
| --- | --- | --- | --- | --- | --- | --- |
|  | x | y | z |  |  |  |
| BLOCK |  |  |  |  |  |  |
| A1 > baseline ∩ T1 > baseline | | | | | | |
| L cerebellum VIII | -24 | -58 | -50 | 334 | 5.23 | 0.025 |
| BEAT |  |  |  |  |  |  |
| A1 > baseline ∩ T1 > baseline | | | | | | |
| L cerebellum VIII | -28 | -56 | -50 | 771 | 5.93 | 0.000 |
| L cerebellum VI | -24 | -54 | -26 |  | 4.15 |  |
| L cerebellum VIIb | -12 | -72 | -42 |  | 4.05 |  |
| R planum temporale | 66 | -34 | 18 | 1065 | 5.67 | 0.000 |
| R planum temporale | 64 | -16 | 12 |  | 3.82 |  |
| R central operculum | 52 | -18 | 16 |  | 4.63 |  |
| R parietal operculum | 44 | -28 | 18 |  | 4.31 |  |
| L parietal operculum | -48 | -32 | 18 | 728 | 5.44 | 0.000 |
| L parietal operculum | -42 | -38 | 22 |  | 5.06 |  |
| L planum temporale | -56 | -28 | 18 |  | 5.14 |  |
| ENVELOPE |  |  |  |  |  |  |
| A1 > baseline ∩ T1 > baseline | | | | | | |
| No suprathreshold clusters |  |  |  |  |  |  |

Notably, auditory beat and envelope recruited slightly different topographies (Fig S1): activations extended more posteriorly (towards the posterior plana temporalia and superior temporal gyri) for auditory beat ([A1 Beat > baseline]) and more anteriorly (towards the anterior superior temporal gyri) for auditory envelope ([A1 Envelope > baseline]). Together, the fMRI results demonstrate that beat information conveyed through audition or touch converges in common temporo-opercular cortices. Conversely, envelope information that goes beyond beat information is conveyed primarily through auditory (but not tactile) signals and activates more anterior regions that extend from plana temporalia towards the anterior superior temporal gyri.

While beat information follows a strictly predictable temporal pattern (in this case, 60 regular pulses per minute), envelope is more variable in time and may therefore demand a longer temporal integration window for accurate prediction and tracking (Hasson et al., 2008, 2015). Using EEG, we therefore directly assessed the temporal evolution and stability of the beat and envelope representations, examining similarities and differences in the auditory (A1) and tactile (T1) conditions.

To assess the neural tracking of beat and envelope information over time, we reconstructed the respective features from EEG activity using a linear ridge-regression model (Crosse, Di Liberto, Bednar, et al., 2016) that was fitted for each sensory modality (A1 or T1) and for each time lag between 0 and 450 ms and comparing them to the respective null-distributions (Fig 3B). In the A1 condition, we found a significant correlation between true and reconstructed beat information for the entire time window (cluster 0-450 ms; p < 0.001) – yet with three distinct peaks at approximately 50 ms, 120 ms and 220 ms. The envelope reconstruction instead revealed a significant correlation a bit later in time, with a cluster between 96 and 177 ms (p = 0.03) followed by a cluster between 290 and 418 ms (p = 0.005). These results nicely dovetail with previous studies showing that while auditory beat modulates neural activity in early time windows (Li et al., 2019), auditory envelope – albeit during speech processing – is tracked over more extended periods (Hausfeld et al., 2018; Puschmann et al., 2019). Interestingly, in the T1 condition we found a significant reconstruction only for beat information (cluster 80-350 ms; p < 0.001), suggesting that touch carries mainly information about the regular temporal structure in music.

To investigate how the brain forms beat and envelope representations over time in either audition or touch, we computed temporal generalization matrices within and across sensory modalities (Fig 3C). The decoding of beat information within the A1 condition showed a significant cluster starting around 65 ms onwards (p < 0.001), with a bigger generalization window that spanned between 150 and 350 ms. Similarly, we found significant decoding of beat information within the T1 condition from 65 ms onwards (p < 0.001), including a similar pattern of temporal generalization. Instead, the decoding of envelope information showed a significant cluster only within the A1 condition. Compared with beat, envelope decoding emerged later in time (between 270 and 350 ms), thereby providing converging evidence that auditory envelope is tracked via sustained representations over more extended periods. Finally, while temporal generalization across sensory modalities did not show any significant results for the envelope features, beat decoding highlighted partially shared patterns of activity across the A1 and T1 conditions. We found significant generalization from T1 to A1 from 176 to 353 ms (p = 0.008) and from A1 to T1 from around 190 to 400 ms (p = 0.048 for cluster 192-241 ms; p = 0.0039 for the cluster 321-401 ms). The generalization from A1 to T1 also highlighted a weak but significant pattern in early time lags from 80 to 96 ms (p = 0.036). Interestingly, this early time window emerged below threshold also in the temporal generalization within the A1 condition.

Collectively, our fMRI and EEG results indicate that both audition and touch convey beat information. While fMRI showed that auditory and tactile beat engage a shared bilateral temporo-opercular cortical system, EEG revealed different underlying temporal dynamics and representations. Auditory beat is tracked instantaneously, whereas tactile beat is reconstructed from later timepoints (after ∼80 ms). Moreover, despite prominent beat decoding from auditory and touch stimulation, we observed only limited and late cross-sensory generalization: shared auditory and tactile beat representations emerge at only later stages (after ∼180 ms). Crucially, based on both fMRI and EEG results, touch is rather poor in contributing envelope information, which is encoded primarily in audition, engaging more anterior temporal regions that extend from plana temporalia towards the anterior superior temporal gyri. Compared with beat, auditory envelope is tracked via sustained representations over later time windows (after ∼270 ms), consistent with the notion that envelope may demand a longer temporal integration window in downstream regions for accurate prediction and tracking (Hasson et al., 2008, 2015).

### Vibrotactile influences on auditory beat and envelope processing in simple scenes

Having established how beat and envelope information is encoded in the auditory and tactile modalities separately, we next asked whether and how this temporal information is integrated into behavioural benefits. In a psychophysics experiment, participants performed a yes-no target detection task with either a unisensory auditory stream (A1), a corresponding unisensory tactile stream (T1) or the combination of the two (A1T1). We then tested whether audiotactile integration improves detection of a brief target (tremolo) embedded in a monophonic (one-stream) music piece by comparing detection sensitivity (d′) and response strategy (*bias*) in the congruent audiotactile condition (A1T1) with the two unisensory conditions (A1, T1; Fig. 4A; Table S1). Observers’ d′ was significantly higher in A1T1 than in A1 (*z* = 2.14, *p* = 0.031, *r* = 0.50) and T1 (*z* = 2.68, *p* = 0.008, *r* = 0.64). Response *bias* was also shifted significantly towards left for A1T1 relative to A1 (*z* = −4.14, *p* < 0.001, *r* = −0.97), and for T1 relative to A1 (*z* = −4.11, *p* < 0.001, *r* = −0.96). In other words, participants were less conservative and biased as well as more sensitive to congruent audiotactile than auditory music. Together, these results show that congruent audiotactile information boosts target detection performance.

**Figure 4.**
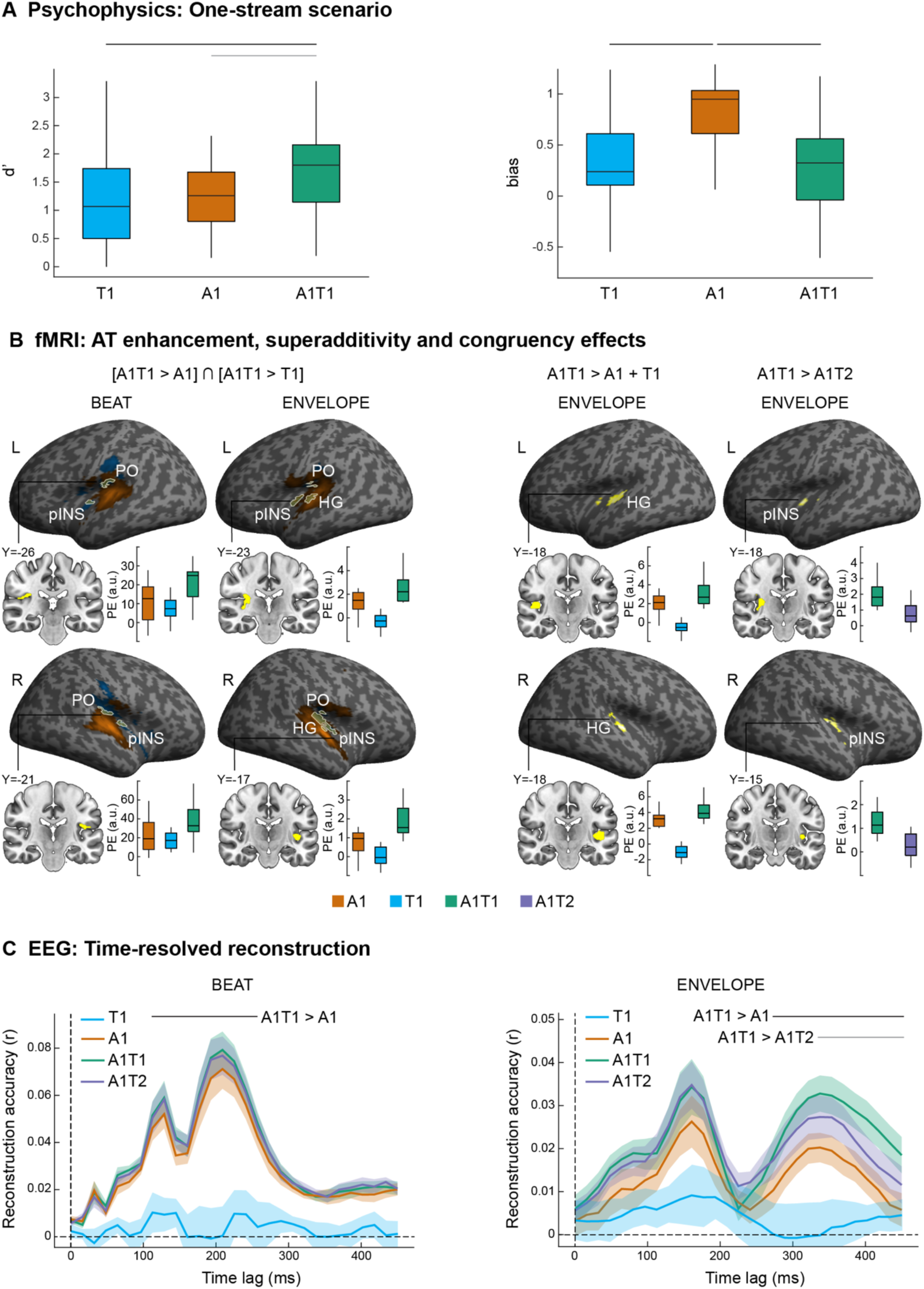
Vibrotactile influences on music processing in simple scenes. **A)** Across-participants’ average d’ (left) and bias (right) for the detection of a target (tremolo) embedded into the music stream. Audiotactile integration increases observers’ d’ and reduces the decisional bias observed for auditory only condition (i.e. observers are more likely to report targets under A1T1 than A1 only condition). Boxplots show median (central line), interquartile range (box), min and max (vertical line). Horizontal lines show statistical significance (black: p < 0.01, grey: p < 0.05). **B**) Left: audiotactile enhancement effect (i.e. [A1T1 > A1] and [A1T1 > T1] for beat (left) and envelope (right). Increased activations for [A1T1 > T1] in orange, [A1T1 > A1] in blue and their overlap in yellow, rendered on an inflated canonical brain (p < 0.001 uncorrected at peak level for visualization purposes, extent threshold k > 0 voxels). White contours represent the formal “conjunction null” conjunction [A1T1 > A1] ∩ [A1T1 > T1]. Representative clusters are also displayed in yellow on coronal slices of a canonical brain. Bar plots show the across-participants’ parameter estimates (PE), averaged across all voxels in the yellow cluster. On the right: superadditivity [A1T1 > A1+T1] and congruency [A1T1 > A1T2] effects for envelope, following the same visualization settings. PO, parietal operculum; HG, Heschl’s gyrus; pINS, posterior insula. A1, one-stream auditory; T1, one-stream tactile; A1T1, one-stream audiotactile congruent; A1T2, one-stream audiotactile incongruent. **C)** Across-participants’ average time-resolved reconstruction of beat (left) and envelope (right) features. Note that reconstruction accuracy is in this analysis based on the decoding model trained on unisensory auditory condition [A1] to examine selectively how auditory representations are influenced by concurrent tactile signals. Plots show mean (line) ± SEM (shaded area). For all plots, reconstruction accuracy is quantified in terms of Fisher-z-transformed Pearson’s correlation (r) between the reconstructed and true feature time-series. We plot results for auditory (A1, orange), audiotactile congruent (A1T1, green) and audiotactile incongruent (A1T2, purple). Horizontal lines show statistically significant temporal clusters (black: p < 0.01, grey: p < 0.05, corrected for multiple comparisons).

Using fMRI, we next investigated whether these audiotactile benefits may rely on audiotactile interactions in primary auditory cortices or in higher-level associative regions (Fig 4B; Table 2). For envelope processing, we observed superadditive interactions (James & Stevenson, 2012; Noppeney, 2012), i.e. greater activations for congruent audiotactile than the sum of unisensory auditory and tactile conditions ([A1T1 > A1 + T1]), in bilateral Heschl gyri, partially extended to the right posterior insula. While we observed a suppressive effect for tactile envelope information alone in primary auditory cortices, it significantly amplified envelope encoding when paired with auditory music. This response profile converges with earlier research showing suppressive effects in primary sensory cortices for signals from non-preferred modalities when presented alone, which turn into BOLD-response amplifications when presented together with signals from the preferred sensory modality (Gau et al., 2020; Leitão et al., 2013; Mozolic et al., 2008; Werner & Noppeney, 2010). This finding suggests that in auditory cortices touch does not furnish its own independent envelope code but sharpens and amplifies auditory temporal representations. A “conjunction null” conjunction [A1T1 > A1] ∩ [A1T1 > T1] analysis further revealed enhanced responses to congruent audiotactile stimulation relative to both unisensory conditions in the right parietal operculum (extending to the rolandic operculum) for beat and envelope processing and bilateral posterior insula for envelope processing. Moreover, the contrast [A1T1 > A1T2] identified stronger activations for audiotactile congruent than incongruent trials in the right posterior insula for envelope processing.

**Table 2.**
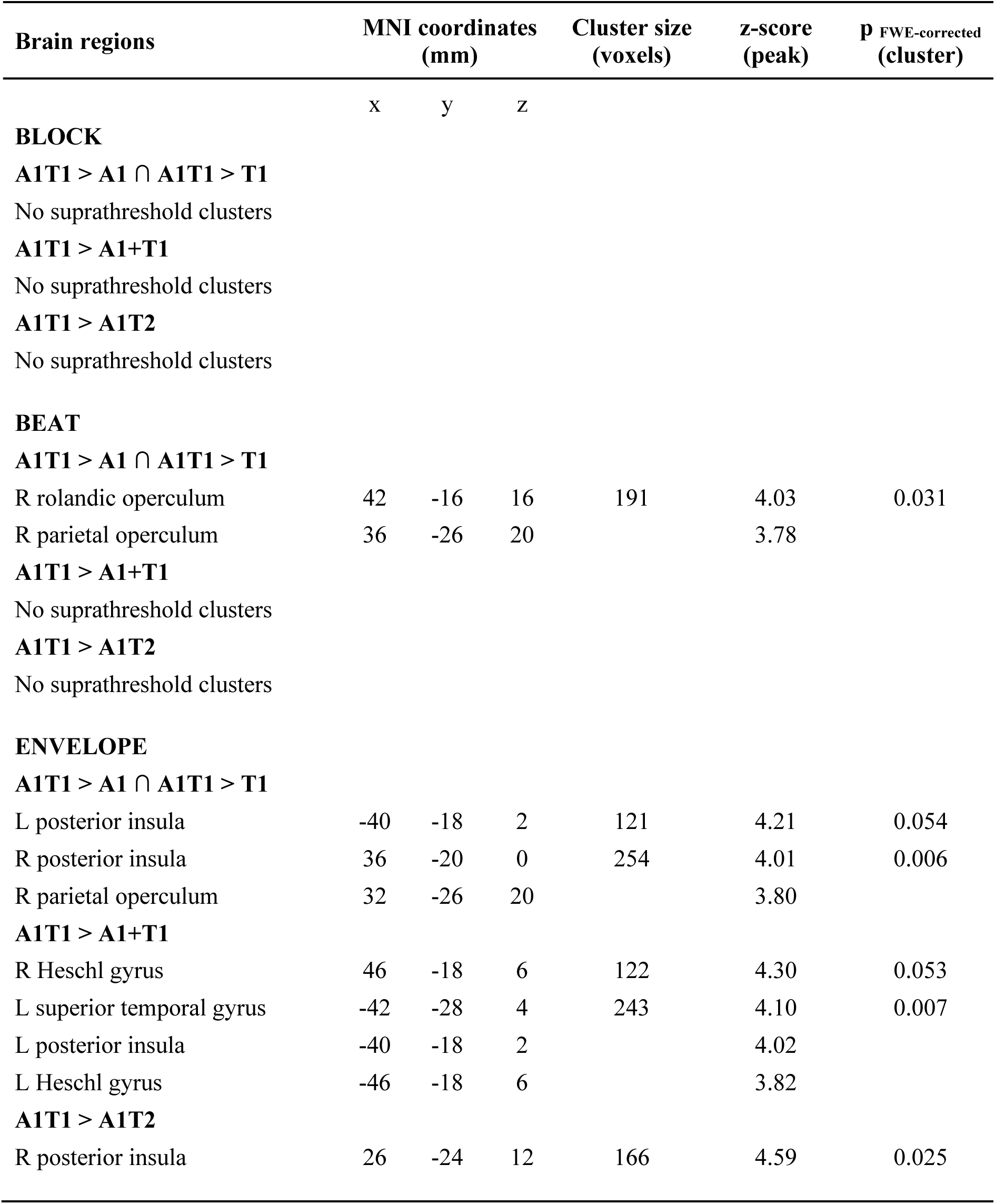
fMRI results: Audiotactile enhancement, superadditivity and congruency. p-values are FWE-corrected at the cluster level for multiple comparisons within the entire brain, with an auxiliary uncorrected peak-level threshold of p < 0.001 (small-volume correction with mask: all > baseline, p < 0.05 FWE peak-level, k > 100). A: auditory; T: tactile; L: left; R: right.

Using EEG and time-resolved stimulus reconstruction analyses, we next characterized whether and how tactile inputs enhanced beat and envelope encoding. We trained a linear ridge-regression model (Crosse, Di Liberto, Bednar, et al., 2016) to reconstruct beat and envelope information from EEG activity under unisensory A1 condition and applied this purely auditory model to EEG data under A1, T1 and A1T1 conditions for each time lag between 0 to 450 ms. Crucially, when applying this A1 model to tactile-only processing, we did not reconstruct beat or envelope information better than chance, demonstrating that the A1 model selectively focuses on auditory beat or envelope information. Therefore, this approach allowed us to assess whether concurrent tactile stimulation enhances the neural encoding of auditory beat and envelope information (Fig 4C). We found a clear temporal dissociation. Touch increased beat reconstruction in an early time window (A1T1 > A1: cluster 112-257 ms; p < 0.001), suggesting that beat information furnished by touch enhances early stages of beat encoding in auditory processing. At later time windows, envelope reconstruction was also significantly better for A1T1 relative to A1 (cluster 273-450 ms; p = 0.003) and for congruent relative to incongruent audiotactile stimulation (A1T1 > A1T2) (cluster 337-450 ms; p = 0.033). Thus, the temporal evolution of the EEG signal correlates more strongly with the audio envelope when touch is present. Moreover, the audiotactile enhancement even exceeded simple linear combination: envelope reconstruction was significantly stronger than the sum of the unisensory auditory and tactile reconstructions (cluster 305-434 ms; p = 0.047; see S2). These later audiotactile effects on envelope encoding align with the notion that the extraction and tracking of envelope information rely on longer temporal integration windows at later and higher-order processing stages along the auditory pathways (Hasson et al., 2008, 2015).

Collectively, our fMRI and EEG findings indicate that concurrent vibrotactile signals enhance the encoding of temporal information in music across multiple processing stages along the cortical hierarchy. Crucially, audiotactile interactions are partly dissociable for periodic temporal structure (beat) and continuous temporal variation (envelope). Beat encoding is amplified at relatively early latencies, with BOLD-response covarying more strongly with beat in the parietal operculum and posterior insula. By contrast, envelope encoding is amplified later in time, with tactile signals supralinearly amplifying envelope representations in primary auditory cortices. This suggests that tactile envelope information sharpens auditory temporal representations rather than generating a general arousal or attention effect.

### Vibrotactile influences on auditory beat and envelope processing in complex scenes

Next, we examined whether vibrotactile signals enhance the segregation and encoding of a temporally correlated auditory stream within more complex polyphonic (i.e. two-stream) scenarios (Fig 5A; Table S1). In a psychophysics experiment, participants performed a yes-no target detection task with a two-stream music piece (A1A2) and a simultaneous vibrotactile stream that was congruent with one of the auditory streams (e.g., A1A2T1). We then evaluated how detection sensitivity (d′) and response strategy (*bias*) changed when the target occurred in the audiotactile congruent stream (A1*A2T1) relative to the incongruent stream (A1A2*T1) and relative to the absence of vibrotactile stimuli (A1A2). We found a significant increase of d′ for A1*A2T1 compared with A1A2*T1 (z = 4.29, p < 0.001, r = 1.00) and A1A2 (z = 2.55, p = 0.011, r = 0.61). Moreover, d′ was significantly higher in A1A2 than A1A2*T1 (z = 4.17, p < 0.001, r = 0.97). Thus, audiotactile congruency significantly improved, whereas incongruency significantly impaired, target detectability relative to unisensory auditory stimulation. In addition, there was a significant leftward shift of *bias* for A1*A2T1 relative to A1A2*T1 (z = -4.29, p < 0.001, r = -1.00) and A1A2 (z = -4.11, p < 0.001, r = -0.98). *Bias* was also significantly leftwards in A1A2 than A1A2*T1 (z = -2.31, p = 0.019, r = -0.54). In other words, audiotactile congruency made participants less conservative, whereas incongruency made them more conservative, relative to unisensory auditory stimulation. Together, these results show that participants’ performance was influenced by a concurrent vibrotactile stream congruent with one of the two auditory streams, suggesting that touch may enhance the segregation and encoding of the temporally correlated auditory information in the brain.

**Figure 5.**
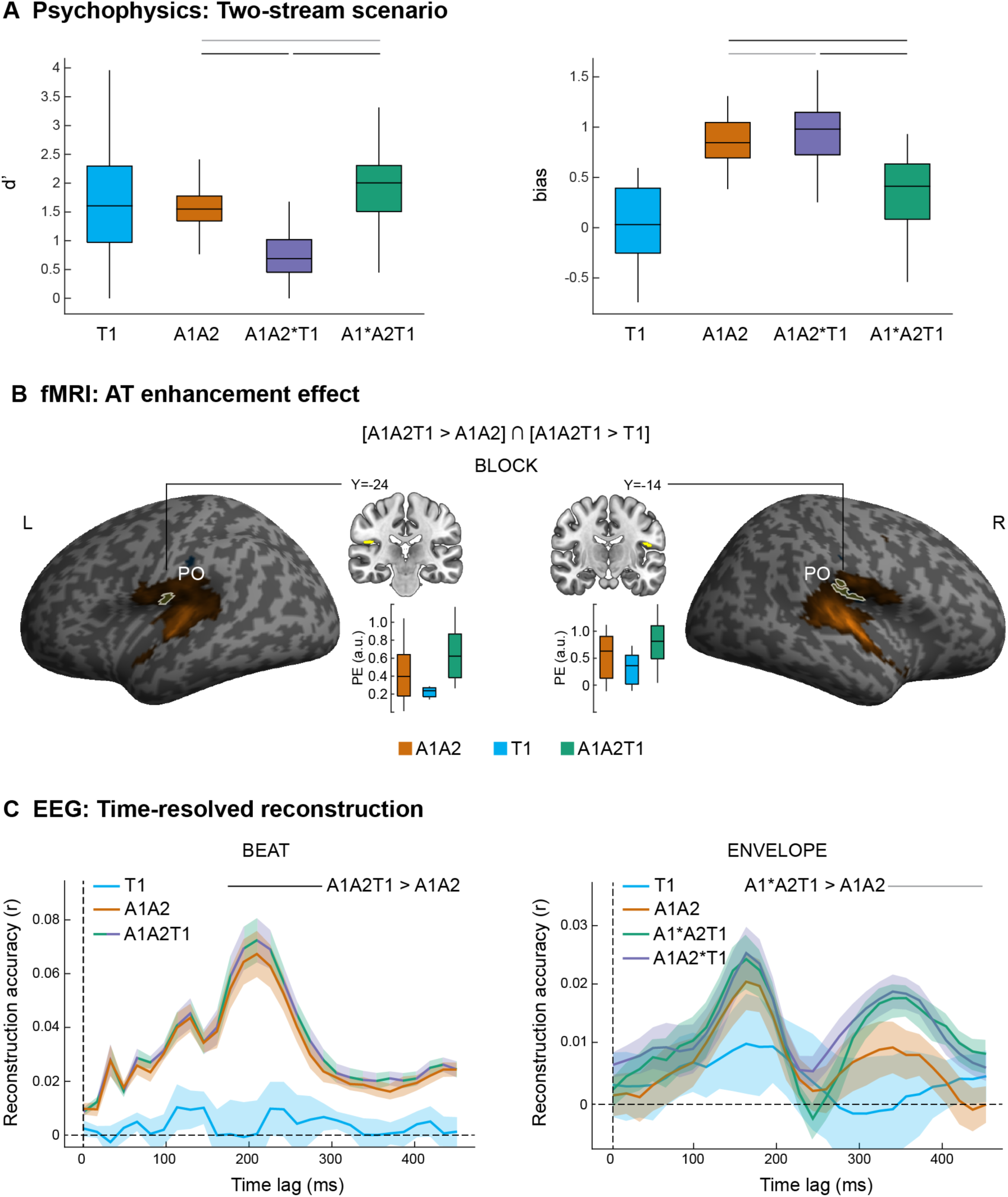
Vibrotactile influences on music processing in complex scenes. **A)** Across-participants’ average d’ (left) and bias (right) for the detection of a target (tremolo) embedded into the music stream. Observers’ d’ increases and the decisional bias decreases when targets are presented in audiotactile congruent streams. Boxplots show median (central line), interquartile range (box), min and max (vertical line). Horizontal lines show statistical significance (black: p < 0.01, grey: p < 0.05). **B)** Audiotactile enhancement effect for the block regressor. Increased activations for [A1A2T1 > T1] in orange, [A1A2T1 > A1A2] in blue and their overlap in yellow, rendered on an inflated canonical brain (p < 0.001 uncorrected at peak level for visualization purposes, extent threshold k > 0 voxels). White contours represent the formal “conjunction null” conjunction [A1A2T1 > A1A2] ∩ [A1A2T1 > T1], which is also displayed as a yellow cluster on coronal slices of a canonical brain. Bar plots show the across-participants’ parameter estimates (PE), averaged across all voxels in the yellow cluster. PO, parietal operculum. A1A2T1, two-stream audiotactile; A1A2, two-stream auditory; T1, one-stream tactile. **C)** Across-participants’ average time-resolved reconstruction of beat (left) and envelope (right) features. In this analysis, the reconstruction accuracy is based on the decoding model trained on the unisensory auditory condition (A1) to examine selectively how auditory representations are influenced by concurrent tactile signals. Plots show mean (line) ± SEM (shaded area). For all plots, reconstruction accuracy is quantified in terms of Fisher-z-transformed Pearson’s correlation (r) between the reconstructed and true feature time-series. For beat, results overlap for auditory (A1A2, orange) and audiotactile (A1A2T1, green/purple) conditions, since beat information is equal across auditory and tactile streams. For envelope, we plot results for auditory (A1A2, orange), audiotactile congruent (A1*A2T1, green) and audiotactile incongruent (A1A2*T1, purple). Horizontal lines show statistically significant temporal clusters (black: p < 0.01, grey: p < 0.05, corrected for multiple comparisons).

Using fMRI, we evaluated how the enhancement driven by congruent tactile stimuli is expressed across the auditory cortical hierarchy. The “conjunction null” conjunction over enhancements of auditory processing by touch and tactile processing by audition [A1A2T1 > A1A2] ∩ [A1A2T1 > T1] revealed activations in the bilateral parietal opercula for the block regressor (p < 0.001 uncorrected peak level), which codes the onset and duration of each condition block, and thus reflects sustained condition-specific activations regardless of beat- and envelope-specific information (Fig 5B). For completeness, when lowering the significance threshold to p < 0.05 uncorrected peak level, we also observed activations for beat processing in bilateral clusters encompassing parietal opercula, posterior insula and the thalamus. Further, we observed superadditive interactions [A1A2T1 > A1A2 + T1] in the bilateral Heschl gyri for envelope processing. It is possible that the presence of two simultaneous auditory streams limited the fMRI ability to detect beat- and envelope-specific effects due to the presence of temporally intermixed stream representations across the auditory hierarchy. Yet, at a lower threshold of significance the activation patterns are qualitatively in line with the one-stream condition: while a higher-level (opercular and posterior insular) cortical system seems responsible for audiotactile enhancement effects, superadditive profiles of audiotactile envelope integration appear in primary auditory cortices.

In EEG, we employed time-resolved reconstruction analyses to compare the encoding strength of beat and envelope across the two-stream conditions with high temporal precision (Fig 5C). Similar to the one-stream scenario, we applied the A1 model trained on A1 stream to EEG data under T1, and the A1 model trained on A1A2 stream to EEG data under A1A2, A1*A2T1 and A1A2*T1, for each time lag between 0 to 450 ms. Importantly, beat was shared across the two auditory streams (A1 and A2) and the tactile stream (T1), thereby providing cues for integrating all three streams. Accordingly, we assessed whether beat information is better represented under audiotactile (A1A2T1) relative to audio-only (A1A2) stimulation. By contrast, envelope differed across the two auditory streams, thereby serving as a cue for stream segregation. Thus, we assessed whether envelope information is better represented for the auditory stream that is congruent with the tactile signal (A1*A2T1) relative to the incongruent auditory stream (A1A2*T1) or any of the two audio-only streams (A1A2). Time-resolved reconstruction was enhanced in the audiotactile relative to the auditory condition, with an earlier enhancement effect for beat information (A1A2T1>A1A2: cluster 170-300 ms; p < 0.001) and a later effect for envelope information (A1*A2T1>A1A2: cluster 321-450 ms, p = 0.009). Similarly to the one-stream condition, the audiotactile enhancement effect surpassed the mere convergence of unisensory auditory and tactile envelope information (Fig S2): despite a complex polyphonic scenario, audiotactile envelope reconstruction was tentatively stronger than the sum of the unisensory auditory and tactile reconstructions (cluster 305-434 ms; p = 0.065). However, the envelope effect did not show stream specificity (A1*A2T1>A1A2*T1, n.s.) and all results emerged in delayed time windows compared with the one-stream scenario, suggesting that a more complex polyphonic scenario may tax the brain’s capability to track congruent time-varying features across the senses. Importantly, results are overall qualitatively consistent across the one-stream and two-stream scenarios: audiotactile beat information is processed in earlier time windows, followed by a later tracking of audiotactile envelope information, possibly due to the need for extended tracking over a longer temporal integration window (Hasson et al., 2008, 2015).

In sum, the fMRI and EEG results obtained in the monophonic (one-stream) conditions generalized to more complex polyphonic (two-stream) scenarios, indicating that congruent vibrotactile signals also support the segregation and amplification of auditory streams amidst distractors in complex audiotactile scenes.

## Discussion

This study offers novel insights into how the brain automatically tracks and merges temporal information from auditory and vibrotactile signals during music perception.

In a psychophysics experiment, we found that congruent vibrotactile stimulation significantly improved the detection of auditory targets embedded in a music stream. Participants showed increased detection sensitivity and a less biased response strategy when exposed to audiotactile congruent stimuli relative to incongruent and audio-only stimuli, in both simple monophonic scenarios and more complex polyphonic scenarios. Together, these findings demonstrate that audiotactile integration boosts the tracking of music information, and suggest that touch may enhance the segregation and encoding of temporally correlated auditory information in the brain (Bizley et al., 2016; Maddox et al., 2015; Shamma et al., 2011; Zion Golumbic et al., 2013).

In subsequent fMRI and EEG experiments, we deliberately chose not to apply a task to focus on more naturalistic, automatic integration. First, we asked whether the brain encodes music information (beat and envelope) from purely auditory and vibrotactile signals via shared neural representations. Time-resolved EEG decoding showed that the brain extracts beat information from both audition and touch in less than 100 ms, complementing previous work on auditory beat processing (Li et al., 2019). Accordingly, touch is potent in conveying the perception of speech rhythm (Guilleminot et al., 2023; Navarra et al., 2014) and musical rhythm (Ammirante et al., 2016; Gilmore & Russo, 2020; Hove et al., 2020; Huang et al., 2012; Large et al., 2015; Tranchant et al., 2017), as also demonstrated by the daily experience of ‘feeling loud music through the body’. Crucially, however, the brain appears to track musical beat via mostly independent neural representations across audition and touch, as suggested by the limited temporal generalization from audition to touch and vice versa. Nevertheless, fMRI regional BOLD-responses showed convergent cortical topographies for auditory and tactile beat processing, suggesting that these dynamic neural representations may evolve in the same cortical networks. Specifically, we found common activations in the bilateral plana temporalia and, interestingly, parietal opercula. The planum temporale is central to music processing (Koelsch, 2011; Koelsch & Siebel, 2005; Peretz & Zatorre, 2005; Vuust et al., 2022), and it has been specifically implicated in beat perception not only through audition (Large et al., 2015) but also through vision and touch (Araneda et al., 2017). The parietal operculum has been traditionally related to the perception of tactile texture (Sathian, 2016) and vibrations (Burton et al., 1993, 2008). However, more recent research has shown that the parietal operculum similarly represents tactile and auditory spectral information (Pérez-Bellido et al., 2018). Our study suggests that planum temporale and parietal operculum may support the extraction of temporal structure, such as beat and envelope information, in both audition and touch. Conversely, we successfully decoded envelope features only in audition and found no temporal generalization across sensory modalities. Similarly, fMRI did not identify common activations across audition and touch for envelope processing. Combined, this suggests that purely vibrotactile signals are rather poor in conveying envelope information, whose extraction may critically depend on more temporally and spectrally fine-grained analyses in the auditory modality (Ding et al., 2014). Importantly, auditory envelope encoding emerged at later time lags than auditory (and tactile) beat encoding. These results nicely dovetail with initial evidence that beat information (transient by nature) modulates neural activity in early time windows (Li et al., 2019), while envelope information (requiring evidence accumulation over time) is tracked over more extended periods (Hausfeld et al., 2018; Puschmann et al., 2019). Collectively, our results indicate that purely auditory and vibrotactile signals successfully convey beat information, while touch per se contributes relatively little to the encoding of envelope information.

Next, we evaluated whether a concurrent vibrotactile signal amplifies the encoding of auditory beat and envelope information in auditory cortices and beyond. Our EEG reconstruction analyses showed tactile enhancement of auditory beat encoding, suggesting that redundant, temporally coherent information across the senses boosts the tracking of naturalistic features (Atilgan et al., 2018; Bizley et al., 2016; Lee & Noppeney, 2011), such as the rhythm of music. Crucially, envelope encoding was more accurate for audiotactile congruent than incongruent and audio-only signals, indicating that the temporal congruency of ongoing audiotactile music enhances the cortical encoding of the music envelope in noise-free conditions (Fu & Riecke, 2023). Thus, multisensory facilitation effects may not necessarily rely on the explicit encoding of purely unisensory information, whose combination strengthens the representation of a common, redundant feature (e.g. beat). Instead, multisensory enhancement might result from non-linear interactions in multisensory neural populations or through cross-feature interactions, where one feature (e.g. envelope) is inferred from a correlated concurrent feature (e.g. beat), as similarly proposed for audiovisual speech envelope tracking (Crosse et al., 2015; Mesgarani et al., 2009). Importantly, the temporal dynamics of audiotactile beat and envelope reconstruction resembled those observed for purely auditory and tactile stimuli: beat encoding emerged earlier than envelope encoding. Thus, our results show that the extraction of music features follows similar temporal dynamics in unisensory and multisensory contexts. Computationally, the brain may leverage temporally aligned and convergent cross-modal information to support naturalistic perceptual scene analysis (Atilgan et al., 2018; Bizley et al., 2016).

While our data intriguingly showed that neural tracking of the music envelope is reduced when auditory and tactile envelopes are incongruent compared to congruent, no significant difference was observed between incongruent and audio-only conditions. Since beat was a redundant signal across the two piano voices (A1 and A2) and across audition and touch, it may have boosted the cortical tracking of envelope for both audiotactile congruent and incongruent streams, for example through cross-feature extraction (Crosse et al., 2015; Mesgarani et al., 2009), thereby reducing possible audiotactile incongruency effects. Importantly, such dampening suggests that the brain was analysing the perceptual scene by considering both beat and envelope cues, with the former enforcing integration (i.e. beat) at earlier time window and the latter segregation (i.e. envelope) at later processing stages. This raises the intriguing hypothesis that the brain solves the causal inference or binding problem in more naturalistic audiotactile scene analysis by combining multiple temporal features operating at different timescales (Noppeney, 2021; Shams & Beierholm, 2022). Building on the present findings, future studies may explicitly manipulate the congruency of the beat and envelope information factorially to investigate their distinct and interactive influences on causal inference.

Interestingly, since participants were passively exposed to the audiotactile stimuli in our fMRI and EEG experiments (with an orthogonal visual task to control for vigilance), our results suggest the automaticity of causal inference despite being exposed to a complex perceptual scene (i.e. polyphonic music). The brain may automatically extract the statistical correlation of consecutive events across the senses, detect and track multiple temporal correspondence cues, which may in turn automatically guide multisensory stream integration versus segregation (Werner & Noppeney, 2010). Rapid beat correspondence may promote initial integration, which is then updated by slower-evolving envelope correspondence cues. A hierarchy of cortical regions may implement such a complex process, as previously demonstrated in the context of Bayesian causal inference with simple, transient stimuli (Aller & Noppeney, 2019; Ferrari & Noppeney, 2021; Mihalik & Noppeney, 2020; Rohe & Noppeney, 2015, 2016). Accordingly, our fMRI results suggest that different types of audiotactile interactions emerged along the cortical hierarchy to support the automatic tracking of music features. On the one hand, we found superadditive profiles of audiotactile envelope integration in primary auditory cortices. In line with fMRI studies on audiovisual integration (Gau et al., 2020; Laurienti et al., 2002; Mozolic et al., 2008; Werner & Noppeney, 2010), superadditivity emerged due to two opposite effects: while vibrotactile input alone led to a deactivation in primary auditory cortices, it amplified the response to a temporally correlated auditory input. Accumulating evidence over the past two decades has shown pronounced cross-modal interactions in early sensory areas (Driver & Noesselt, 2008; Ghazanfar & Schroeder, 2006; Kayser & Logothetis, 2007; Noppeney, 2021; Schroeder & Foxe, 2005). Notably, there is electrophysiological evidence of superadditive profiles of audiotactile integration in the caudal belt of non-human primates (Kayser et al., 2005, 2008; Schroeder et al., 2008). Moreover, lip movements enhance the EEG tracking of a congruent speech stream, possibly in human auditory cortex (Crosse et al., 2015). Together, our fMRI results strengthen the present literature by showing automatic audiotactile integration in the primary human auditory cortex during naturalistic music listening. Conversely, a higher-level opercular and posterior insular system was associated with audiotactile enhancement and congruency effects; further, converging evidence from EEG time-resolved reconstruction showed that congruency effects emerged at later time lags (> 300 ms). Thus, recognizing audiotactile incongruency might be implemented down the hierarchy, thereby supporting the arbitration between audiotactile integration and segregation (i.e. causal inference).

Finally, we examined whether a simultaneous vibrotactile signal enhances the segregation and encoding of auditory streams in more complex multi-stream scenarios. Time-resolved EEG reconstruction results from the two-stream conditions were in line with the one-stream conditions: beat encoding was more accurate for audiotactile than audio-only streams and envelope encoding was more accurate for audiotactile congruent than incongruent and audio-only streams. However, results did not show improved envelope encoding for audiotactile congruent than incongruent streams. Importantly, the temporal dynamics of beat and envelope reconstruction resembled those observed for the one-stream condition: audiotactile enhancement effects appeared at earlier time lags for beat than envelope encoding, suggesting that in general envelope information may demand a longer temporal integration window in downstream regions for accurate prediction and tracking (Hasson et al., 2008, 2015). Interestingly, however, both the beat and envelope enhancement effects were delayed in the two-stream scenario compared to the one-stream scenario. This suggests that the brain may require additional time to extract multisensory features embedded in a more complex multi-stream environment. The need to parse multiple concurrent information streams across the senses may, in fact, strain our capacity-limited attentional resources (Alsius et al., 2005; Talsma et al., 2010; Tang et al., 2016), resulting in a delay—a hypothesis aligning with the idea that attention accelerates cortical tracking (Riecke et al., 2019). Despite these differences in temporal dynamics, fMRI topographies were qualitatively similar across the one-stream and two-stream conditions: while an opercular and posterior insular system was associated with audiotactile enhancement effects, superadditive profiles of audiotactile envelope integration appeared predominantly in primary auditory cortices. Notably, beat processing did not elicit superadditivity effects in either the one-stream or two-stream condition. This suggests that beat information may already be maximally tracked in audition, leaving little room for additional superadditive enhancement due to a ceiling effect (James & Stevenson, 2012; Noppeney, 2012).

Together, our results indicate that congruent vibrotactile signals also support the segregation and amplification of auditory streams amidst distractors. This evidence resonates with electrophysiological work in audiovisual processing showing that lip movements facilitate the tracking of speech in cocktail-party scenarios (Crosse, Di Liberto, & Lalor, 2016; Zion Golumbic et al., 2013), possibly leveraging the statistical temporal coherence between the concurrent visual and auditory streams (Atilgan et al., 2018; Lee & Noppeney, 2011). The automatic grouping of coherent cross-modal features into a multisensory perceptual unit may promote its salience and attentional selection, and thereby, its cortical tracking despite the presence of concurrent distractors (Bizley et al., 2016; Maddox et al., 2015; Shamma et al., 2011; Zion Golumbic et al., 2013). The same core computational mechanism may be at play when processing audiotactile pairings: the congruency among audition and touch may support stream segregation and thereby guide auditory scene analysis by leveraging temporally aligned and convergent audiotactile information despite a complex multi-stream scenario. Accordingly, there is mounting behavioural evidence that envelope-modulated vibrotactile streams boost speech-in-noise understanding (Cieśla et al., 2022; Fletcher et al., 2024; Răutu et al., 2023; Schulte et al., 2023).

In conclusion, our results show that touch clearly conveys beat information but is rather poor in contributing envelope information. Yet, both beat and tactile envelope information modulate music processing across the auditory cortical hierarchy to support auditory scene analysis in the human brain.

## Methods

### Ethics statement

All volunteers provided written informed consent and received financial reimbursement or course credits for their participation in the experiment. The study was approved by the ethical review committee of the University of Birmingham (approval number: ERN_15-1458P) and was conducted following the principles outlined in the Declaration of Helsinki.

### Participants

A first group of participants underwent the psychophysics experiment in one day. A total of 26 volunteers participated in the experiment. Two of those volunteers were excluded based on a priori criteria (see section “Inclusion criterion: congruency judgement”). As a result, 24 participants (3 males; mean age 22, range 18 to 30 years) were included in the analysis and results of the psychophysics experiment. This final number of included participants was based on an a priori power analysis (G*Power 3.1; Faul et al., 2009), with power (1-β) = 0.8, α = 0.05 and effect size Cohen’s d_AV_ = 0.6. Estimation of effect size was derived from a two-tailed paired sample t-test evaluating the effect of touch (one-stream: A1T1 vs A1; two-stream: A1*A2T1 vs A1A2*T1) on target detection (d′) in a preliminary pilot study (N=9).

A second group of participants completed the fMRI and EEG experiments over two weeks. Twelve participants (3 males; mean age 28 years, range 22-34 years) took part in the experiments and were included in the analyses and results based on a priori criteria (see section “Inclusion criterion: congruency judgement”). To maximize within-participant precision, we included four hours of data acquisition separately for each imaging modality (see similar sample sizes and analysis approaches: Aller et al., 2022; Aller & Noppeney, 2019; Alluri et al., 2012, 2013; Ferrari & Noppeney, 2021; Hoefle et al., 2018; Mihalik & Noppeney, 2020; Sankaran et al., 2018; Santoro et al., 2017; Toiviainen et al., 2014).

All volunteers reported normal or corrected to normal vision, normal hearing and touch and no history of neurological or psychiatric conditions; they were right-handed according to the Edinburgh Handedness Inventory (Oldfield, 1971; mean laterality index: 85; range: 60–100); they had never received any formal music training and were classified as non-musicians via the Music USE (MUSE) Questionnaire (Chin & Rickard, 2012) based on duration, frequency and regularity of instrument playing.

### Stimuli

Auditory stimuli consisted of counterpoint music pieces custom-made by a professional composer and synthesized from Musical Instrument Digital Interface (MIDI) files using Linux MultiMedia Studio 1.1.3 (LMMS) with a piano sound font (grand-piano-YDP-20160804). Tactile stimuli corresponding to these auditory piano pieces were synthesized using the LMMS TripleOscillator. We first extracted the three lowest harmonics of each note. To respect the vibratory limits of the piezoelectric presentation device, we set the three sinusoidal carrier frequencies 1 octave lower than the original score. These carrier frequencies were thus in the range 60 and 400 Hz, known to drive Pacinian mechanoreceptors (Kandel et al., 2000). Next, we extracted the auditory envelope for each music piece by computing the root-mean-squared (RMS) amplitude of each piano sound. We applied a lowpass moving average filter with a 20 ms window. The three sinusoidal carrier frequencies were modulated with these lowpass filtered auditory envelopes. As a result, we obtained auditory and tactile stimuli that shared beat (60 bpm) and envelope information for each music piece (Fig 1A). Auditory and tactile synthesized stimuli were recorded at 44100 Hz with a 16-bit resolution, normalized and saved as WAV files using Audacity 2.1.2.

We created 24 different counterpoint music pieces based on the scores of the professional composer (duration: 8 s in psychophysics, 28 s in fMRI and EEG; mean sound pressure level: 75 dB). For the two-stream conditions, two auditory streams were combined into two-stream polyphonic pieces via simultaneous recording according to the composer’s score. To generate audiotactile incongruent stimuli, we selected pairs of envelopes with minimal temporal correlations from a set of 10,000 possible permutations. For each music piece, we selected the pair that gave the lowest overall temporal correlation score (one-stream condition: 0.33 ± 0.01, two-stream condition: 0.32 ± 0.02, across-streams’ mean ± SEM; see Fig 1B).

In the psychophysics experiment, participants performed a yes-no target detection task. The target consisted of a 2 Hz sinusoidal modulation of envelope intensity called “tremolo”, which was inserted 300 ms after the onset of a note using Audacity (1700 ms duration, 100 ms fade-out).

### Inclusion criterion: congruency judgement

In a prior psychophysics screening (Fig 1C), volunteers were pre-selected based on the ability to perceptually discriminate between audiotactile congruent and incongruent stimuli (duration: 8 s; mean sound pressure level: 75 dB). To avoid interference with the main experiments, the screening included a distinct set of 15 congruent and 15 incongruent audiotactile music pieces. In each trial, observers reported whether the audio and tactile streams were temporally congruent or incongruent via pedal press (yes: left pedal; no: right pedal). The experimental setup was identical to the psychophysics and EEG experiments (see section “Experimental setup”). Each participant completed 2 experimental runs, i.e. 2 conditions (AT congruent vs. incongruent) × 15 trials / condition / run × 2 runs = 60 trials in total. Based on signal detection theory (Wickens, 2002), “yes” responses to congruent and incongruent trials were classified as hits and false alarms, respectively. For each participant, we then calculated the following indices:

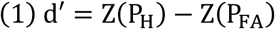

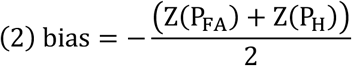

with P_H_ = proportion of hits, P_FA_= proportion of false alarms. Volunteers who showed d′ > 2 (threshold defined in a preliminary pilot with 9 participants) were included in the study.

### Experimental design and procedure

In the psychophysics experiment, participants reported whether a target (“tremolo”, see section “Stimuli”) was present in the music stream in a yes-no target detection task. The experiment comprised six conditions (Fig 2A): one-stream auditory (A1); one-stream tactile (T1); one-stream audiotactile congruent (A1T1); two-stream auditory (A1A2); two-stream audiotactile with target in the congruent auditory and tactile streams (A1*A2T1) or in the incongruent auditory stream only (A1A2*T1). For each condition, targets appeared in 50% of the trials. For each participant, stream identity (stream 1 / stream 2) and target position (first / second half of the stream) were counterbalanced across conditions to minimize any target onset expectations. Participants were instructed to concentrate on the auditory and tactile stimuli with their eyes closed. After stimulus presentation (8 s), they reported as accurately as possible whether a target was present via pedal press (yes: left pedal; no: right pedal). Each new trial started after the participant’s response. Participants firstly completed 5 experimental runs for the one-stream scenario, in which we presented 3 (T1, A1, A1T1) × 2 (target, no target) conditions (6 conditions × 8 trials / condition / run × 5 runs = 240 trials in total). Participants then completed 7 experimental runs for the two-stream scenario, in which we presented 4 (T1, A1A2, A1*A2T1, A1A2*T1) × 2 (target, no target) conditions. Notice, however, that A1*A2T1 and A1A2*T1 shared the same “no target” trials, for a total of 7 conditions (7 conditions × 6 trials / condition / run × 7 runs = 294 trials in total). Trials were presented in random order for each participant and run. Participants familiarized themselves with the stimuli and procedure via one preliminary practice run at the beginning of the one-stream and two-stream runs, respectively.

The fMRI and EEG experiments included six experimental conditions (Fig 2B): one-stream auditory (A1), one-stream tactile (T1), one-stream audiotactile congruent (A1T1), one-stream audiotactile incongruent (A1T2), two-stream auditory (A1A2) and two-stream audiotactile (A1A2T1). In the two-stream audiotactile (A1A2T1) condition, the tactile envelope was congruent with one of the two auditory streams in the polyphonic piece and incongruent with the other stream (e.g. T congruent to A1 is denoted as A1*A2T1). Each condition was presented separately in a stimulation block (duration: 28 s, i.e. presentation of one music piece per block) followed by a baseline block (duration: 6 s in fMRI, 2 s in EEG). Participants were instructed to concentrate on the auditory and tactile stimuli with their eyes closed. To maintain attention and avoid response- and task-related confounds, participants reported the rare occurrences of brief full-screen light-grey flashes (luminance: 85 cd/m^2^; duration: 50 ms) interspersed during the stimulation blocks via pedal press with their right foot. Luminance was adjusted to optimize flashes’ visibility with eyes closed. For the fMRI experiment, every participant completed 16 runs over two days, with 18 stimulation blocks per run (3 blocks / condition / run × 6 conditions × 16 runs = 288 blocks in total). For the EEG experiment, every participant completed 12 runs over two days, with 24 stimulation blocks per run (4 blocks / condition / run × 6 conditions × 12 runs = 288 blocks in total). The order of stimulation blocks was pseudo-randomized within participants to avoid successive repetitions of the same condition and counterbalance the order of conditions across experimental runs. Participants familiarized themselves with the stimuli and procedure via one preliminary practice run at the beginning of each neuroimaging day (both for fMRI and EEG). In each run, 6 stimulation blocks contained flashes (3 blocks with 1 flash; 3 blocks with 2 flashes). We randomized flashes’ onsets with the following constraints: no flashes within the first and last 2 seconds of a block; a minimum gap of 2 seconds between two consecutive flashes. The number of flashes was counterbalanced across conditions within each participant.

### Experimental setup

The experiment was presented via Psychtoolbox (Kleiner et al., 2007) version 3.0.15 running under MATLAB R2018b (MathWorks Inc.) on a Linux machine (Ubuntu 18.04.2 LTS). Stimuli were extracted from WAV files and played via MATLAB custom-code. Tactile stimuli were presented with a piezoelectric device (PTS-C2, Dancer Design, UK) via an external sound card (Asus Xonar U7, Taiwan). A piezoelectric stimulation head was applied to the fingertip of each index finger.

For the psychophysics and EEG experiments, auditory stimuli were presented with EEG-compatible earplugs (EARTONE, Insert Earphone 3A) controlled via the stimulation PC’s built-in soundcard. Visual stimuli were presented via an LCD monitor (30” Dell UltraSharp U3014, USA; 646 2560 × 1600 pixels resolution; 60 Hz frame rate) at a viewing distance of ∼60 cm. Participants were instructed to sit still in a dimly lit cubicle with their eyes closed and their head positioned on a chin rest. They gave responses by pressing a pedal with their right foot (SODIAL, Shenzhen IMC Digital Technology Co.). Stimuli presentation was accompanied by background white noise presented with the headphones (65 dB sound pressure level) to mask the sound of tactile vibrations.

For the fMRI experiment, auditory stimuli were presented with an MR-compatible system (SOUNDPixx MRI pneumatic transducer and amplifier VPX-ACC-8100, QC Canada; MRIaudio in-ear headphones, USA) controlled via the stimulation PC’s built-in soundcard. Visual stimuli were back-projected onto a Plexiglas screen using a Barco Present-C F-Series projector (F35 WUXGA, UK; 1920 × 1024 pixels resolution; 60 Hz frame rate) and they were visible to the participants via a mirror mounted on the MR head-coil at a viewing distance of ∼68 cm. Participants were instructed to lay still in the scanner with their eyes closed. They gave responses by pressing an MR-compatible keypad (NATA LXPAD 1×5-10M, BC Canada) attached to the right foot with an elastic cohesive bandage and secured via foam supports.

### Psychophysics data analysis

Based on signal detection theory (Wickens, 2002), “yes” responses in target and no target trials were classified as hits and false alarms, respectively. For each participant and experimental condition, we then calculated d’ prime and bias using standard formulae (see section ‘Inclusion criterion: congruency judgment’). After rejection of normality (Kolmogorov-Smirnov Test), individual d′ and bias were entered into group pair-wise comparisons across conditions via two-tailed non-parametric Wilcoxon signed-rank tests.

### Music feature extraction

The beat information was modelled as a Dirac comb (i.e. an impulse train of delta functions). We applied a periodic function with a fixed period of T= 1s, which corresponds to the 60bpm at which the MIDI files were recorded:

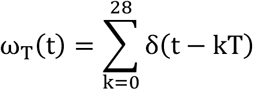

Using the WAV audio files, the envelope information was computed by first band-pass filtering each music piece with a filter bank of 8 logarithmically spaced filters between 100-8000 Hz. We then calculated the Hilbert transform of each signal obtained from the filter bank. The final envelope was obtained by averaging the 8 resulting analytic signals (Yang et al., 1992).

### MRI data acquisition and analysis

#### Data acquisition

A 3T Siemens Prisma MR scanner was used to acquire both a T1-weighted anatomical image (TR = 2000 ms, TE = 2.03 ms, flip angle = 8°, FOV = 256 mm × 256 mm, 208 sagittal slices acquired in sequential ascending direction, voxel size = 1 × 1 × 1 mm^3^) and T2*-weighted axial echoplanar images (EPI) with blood-oxygenation-level-dependent contrast (gradient echo, multiband factor of 2, TR = 1550 ms, TE = 35 ms, flip angle = 71°, FOV = 210 × 210 × 150 mm^2^, 60 axial slices acquired in interleaved ascending direction, voxel size = 2.5 × 2.5 × 2.5 mm^3^, no interslice gap). For each participant, a total of 400 volumes × 16 runs were acquired, after discarding the first four volumes of each run to allow for T1 equilibration effects. Data acquisition was performed over 2 days. The anatomical image was acquired at the end of day 1.

#### Preprocessing

The data were preprocessed and analyzed using Statistical Parametric Mapping (Friston et al., 1994) (SPM12; fil.ion.ucl.ac.uk/spm). Scans from each participant were realigned using the first scan as a reference, unwarped, spatially normalized into MNI standard space using parameters from the segmentation of the T1 structural image, spatially smoothed with a Gaussian kernel of 8 mm full-width at half-maximum, and resampled to a spatial resolution of 2 × 2 × 2 mm^3^. The time series of all voxels were high-pass filtered to 1/128 Hz.

#### General linear model specification

The fMRI experiment was modelled as a mixed block-event related design with each regressor entered in the design matrix after being convolved with the canonical hemodynamic response function. We modelled the 6 experimental conditions (one-stream auditory: A1, one-stream tactile: T1, one-stream audiotactile congruent: A1TI, one-stream audiotactile incongruent: A1T2, two-stream auditory: A1A2, two-stream audiotactile: A1A2T1) as blocks. Each block was modulated by beat and envelope information as separate parametric modulators. Importantly, the A1A2T1 condition included three parametric modulators: one parametric modulator for the beat information that is shared across the two auditory and the tactile streams and two parametric modulators for the envelope information, i.e. one encoding the envelope of melody 1 and one encoding the envelope of melody 2 in the counterpoint music piece. Please note that parametric modulators for the envelope information have been orthogonalized with respect to the parametric modulators for the beat information. Hence, they reflect on the variance that is orthogonal to the beat information. Onsets of all flashes were modelled as events (duration: 0 s) as a separate condition of no interest. Nuisance covariates included the six realignment parameters to account for residual motion artefacts. Condition-specific effects (i.e. parameter estimates for the canonical hemodynamic response function regressors) for each subject were estimated according to the general linear model and passed to a second-level analysis as contrasts. This involved creating 6 contrast images that corresponded to experimental blocks (A1, T1, A1T1, A1T2, A1A2, A1A2T1), 6 contrast images that corresponded to beat parametric modulators (A1, T1, A1T1, A1T2, A1A2, A1A2T1) and 7 contrast images that corresponded to envelope parametric modulators (A1, T1, A1T1, A1T2, A1A2, A1*A2T1, A1A2*T1). The resulting 19 contrast images (i.e., each regressor of interest relative to the unmodelled fixation condition, summed over the 16 runs) for each subject were entered into a second-level ANOVA. Inferences were made at the second level to allow for a random-effects analysis and inferences at the population level.

#### Statistical comparisons

Our analysis focused on three aspects:

##### 1. Encoding beat and envelope information from audition and touch

We investigated the encoding of beat (respectively, envelope) information in audition and touch by comparing the beat and the envelope parametric modulators against baseline separately for each sensory modality. To identify audiotactile convergence regions for beat or envelope information we specified a “conjunction null” conjunction [A1 > baseline] ∩ [T1 > baseline].

##### 2. Vibrotactile influences on auditory beat and envelope processing in simple scenes

We investigated how the brain combines beat (respectively, envelope) information from audition and touch using three approaches: i. the “conjunction null” conjunction [A1T1 > A1] ∩ [A1T1 > T1] identified enhancement for audiotactile stimulation relative to unisensory conditions common to auditory and tactile stimulation; ii. the contrast [A1T1 > A1 + T1] identified superadditive interactions as an index for audiotactile integration; iii. the contrast [A1T1 > A1T2] identified stronger activations for audiotactile congruent than incongruent stimuli.

##### 3. Vibrotactile influences on auditory beat and envelope processing in complex scenes

We examined whether a vibrotactile signal modulates auditory processing in more complex scenes with two competing auditory streams in terms of the average activation (block regressor) and specifically for envelope or beat information: i. the “conjunction null” conjunction [A1A2T1 > A1A2] ∩ [A1A2T1 > T1] identified the enhancement of audiotactile stimulation relative to unisensory conditions common to auditory and tactile stimulation; ii. the contrast [A1A2T1 > A1A2 + T1] identified superadditive audiotactile interactions. For the envelope analyses, we specified these contrasts selectively for the auditory envelope that is congruent to the tactile envelope: i. the “conjunction null” conjunction [A1*A2T1 > A1A2] ∩ [A1*A2T1 > T1] identified the enhancement of audiotactile stimulation relative to unisensory conditions common to auditory and tactile stimulation; ii. the contrast [A1*A2T1 > A1*A2 + T1] identified superadditive interactions.

Unless otherwise stated, we report activations family-wise error (FWE)-corrected for multiple comparisons at p < 0.05 cluster level with an auxiliary uncorrected peak-level threshold of p < 0.001. We correct for multiple comparisons within a search volume defined by [audiotactile stimulation > baseline] at p < 0.05 FWE (peak-level).

### EEG data acquisition and analysis

#### Data acquisition

Continuous EEG signals were recorded from 64 channels using AgAgCl active electrodes arranged in a 10/20 layout, referenced at FCz (ActiCapSlim, Brain Products GmbH, Gilching, Germany). Signals were digitized at 5000 Hz with an anti-aliasing filter of 1000 Hz, then down-sampled to 1000 Hz with a high-pass filter of 0.1 Hz and low-pass filter of 500 Hz. Electrode impedances were kept below 20 kOhm. Triggers from the stimulus-control computer were sent via LabJack to the EEG acquisition computer. Data acquisition was performed over 2 days.

#### Preprocessing

Preprocessing was performed with the FieldTrip toolbox (Oostenveld et al., 2011) (fieldtriptoolbox.org). Raw data were high-pass filtered at 0.3 Hz, low-pass filtered at 150 Hz and band-stop filtered around the line noise and its harmonics (49-51 Hz, 99-101 Hz, and 149-151 Hz). Data were epoched for each music piece from 0 s to 28 s. Trials were subsequently visually inspected for artefacts. Independent component analysis was used to assess artefacts due to eye movement. Given that participants kept their eyes closed throughout the experiment, we expected only negligible eye movements. Indeed, we did not remove any components due to eye movements for any participants. Finally, the EEG recording was re-referenced using the two mastoids (TP9 and TP10), low-pass filtered to 32 Hz and down-sampled to 64 Hz.

#### Time-resolved beat and envelope reconstruction

To investigate how the brain tracks and encodes beat and envelope information in the six different experimental conditions we used single-lag (i.e. time-resolved) multivariate stimulus reconstruction (Crosse, Di Liberto, Bednar, et al., 2016) based on 64 channel EEG signals. To assess the contribution of EEG signals at individual time lag to the stimulus reconstruction and thereby provide insights into the temporal evolution of beat and envelope information in the brain, we trained the decoders on EEG signals at individual lags within a time window from 0 to 450 ms. For illustration, the matrix R for the single lag of 100ms and 64 EEG channels then becomes:

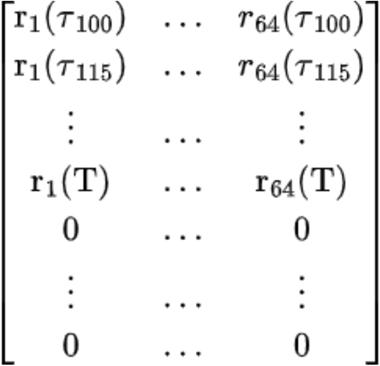

#### Temporal generalization

We characterized the stability and dynamics of beat and envelope representations extracted from audition or touch using the temporal generalization method (King & Dehaene, 2014). Here, the time-resolved response functions were trained at time point t to learn the mapping from, e.g., EEG signals for unisensory sensory touch or audition to the music’s beat (respectively, envelope). This learnt mapping was then used to predict the beat (respectively, envelope) features from the unisensory tactile (or auditory) EEG activity patterns across all other time points. The Pearson’s correlation coefficient between the true and decoded stimulus feature was computed and entered into a training time × generalization time matrix. We computed training time × generalization time matrix within each sensory modality (e.g. from unisensory auditory to unisensory auditory conditions) and between sensory modalities (e.g. from unisensory auditory to unisensory tactile).

#### Statistical comparisons

Our analysis focused on three aspects:

##### 1. Encoding beat and envelope information from audition and touch

The single lag decoding models were trained and tested on the same and opposite sensory modalities. Within the limits imposed by the spatial resolution of EEG, we used temporal generalization matrices to assess the stability and dynamics of the beat and envelope representations within each modality (King & Dehaene, 2014). Next, we assessed temporal generalization across sensory modalities to provide initial insights into the similarity and distinctiveness of neural generators and representations involved in coding beat and envelope information across audition and touch. While successful temporal generalization must be interpreted with caution given the low spatial resolution of EEG, the absence of cross-sensory generalization suggests that beat, respectively envelope, representations rely on modality-specific coding strategies.

##### 2. Vibrotactile influences on auditory beat and envelope processing in simple scenes

To investigate whether a concurrent vibrotactile signal can amplify the neural encoding of auditory beat and envelope information, we used the time-resolved decoding models for beat (respectively, envelope) trained on the one-stream auditory condition (i.e. A1) and generalized them to the audiotactile congruent (A1T1) and incongruent (A1T2) conditions, and compared the reconstruction accuracy across conditions. In cases where auditory and tactile beat (respectively, envelope) information can be shown to rely on separate representations, training the time-resolved decoding models selectively on A1 ensures that we selectively assess whether vibrotactile signals amplify the neural encoding of auditory beat or envelope representations.

##### 3. Vibrotactile influences on auditory beat and envelope processing in complex scenes

Similarly, we generalized the single lag decoding models trained on one stream of the A1A2 condition to the two audiotactile conditions (A1*A2T1, A1A2*T1) and compared the reconstruction accuracies across conditions. Because the beat is shared across all music pieces, we investigated whether a concurrent vibrotactile signal that is temporally correlated with one of the two auditory streams modulates the encoding of auditory beat information. For the envelope information, we specifically examined whether a vibrotactile signal enhances the segregation and neural encoding of the temporally correlated auditory streams by directly comparing the reconstruction accuracy of audiotactile congruent (A1*A2T1) and audiotactile incongruent (A1A2*T1) envelope.

#### EEG statistics

For the time-resolved reconstruction and temporal generalization analyses, we entered the subject-specific time courses of Fisher-z-transformed Pearson’s correlation coefficients into a Monte-Carlo cluster-based permutation test using a one-sample t-statistic and 4096 permutations (Nichols & Holmes, 2002). For the envelope reconstruction, the null distribution was generated by randomly permuting, within each subject, the assignment between EEG trials (i.e. blocks) and their corresponding stimulus envelopes, thereby disrupting the true pairing while preserving the temporal autocorrelation of each signal. For the beat reconstruction, permuting trials would not generate a null distribution because all music stimuli had the same beat. Instead, we generated the null distribution by circularly shifting the beat time course relative to the EEG signal by a random temporal offset within each subject and permutation, thereby destroying the temporal alignment but preserving the temporal autocorrelation. We performed multiple-comparison correction across one-dimensional (time), or two-dimensional (time × time) data based on cluster-level inference by computing the maximum summed t-value within each cluster with an auxiliary cluster-defining threshold of p < 0.05 (uncorrected). Unless otherwise stated, we report clusters at two-tailed p < 0.05, cluster-level corrected.

## Data and code availability

The data and materials necessary to replicate the current findings are available on the Open Science Framework and GitHub (links provided to reviewers); upon publication, a persistent identifier (DOI) will be made publicly available.

## Acknowledgments

This research was funded by the European Research Council (ERC Starting Grant Multsens 309349; ERC Advanced Grant MakingSense 101096659). The authors thank Scott Rubin for composing the music pieces.

