## Supplementary files for "Feeling the music: Audiotactile encoding of temporal structure in the human brain"

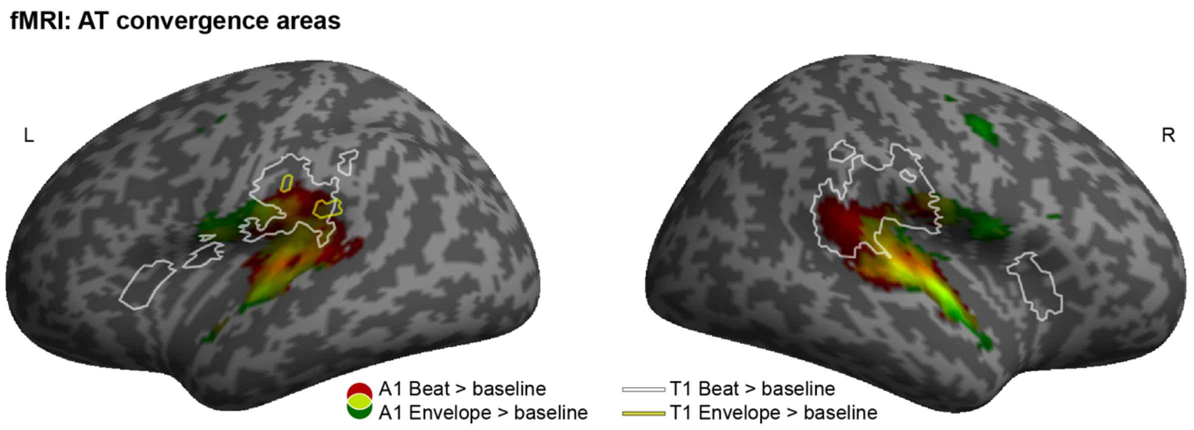


**Figure S1 | Audiotactile convergence: comparison between beat and envelope.** Increased activations for [A1 Beat > baseline] in red, [A1 Envelope > baseline] in green and their overlap in yellow, rendered on an inflated canonical brain ($p < 0.001$ uncorrected at peak level for visualization purposes, extent threshold $k > 0$ voxels). Additionally, white contours represent increased activations for [T1 Beat > baseline] and yellow contours represent increased activations for [T1 Envelope > baseline]. A1, one-stream auditory; T1, one-stream tactile.

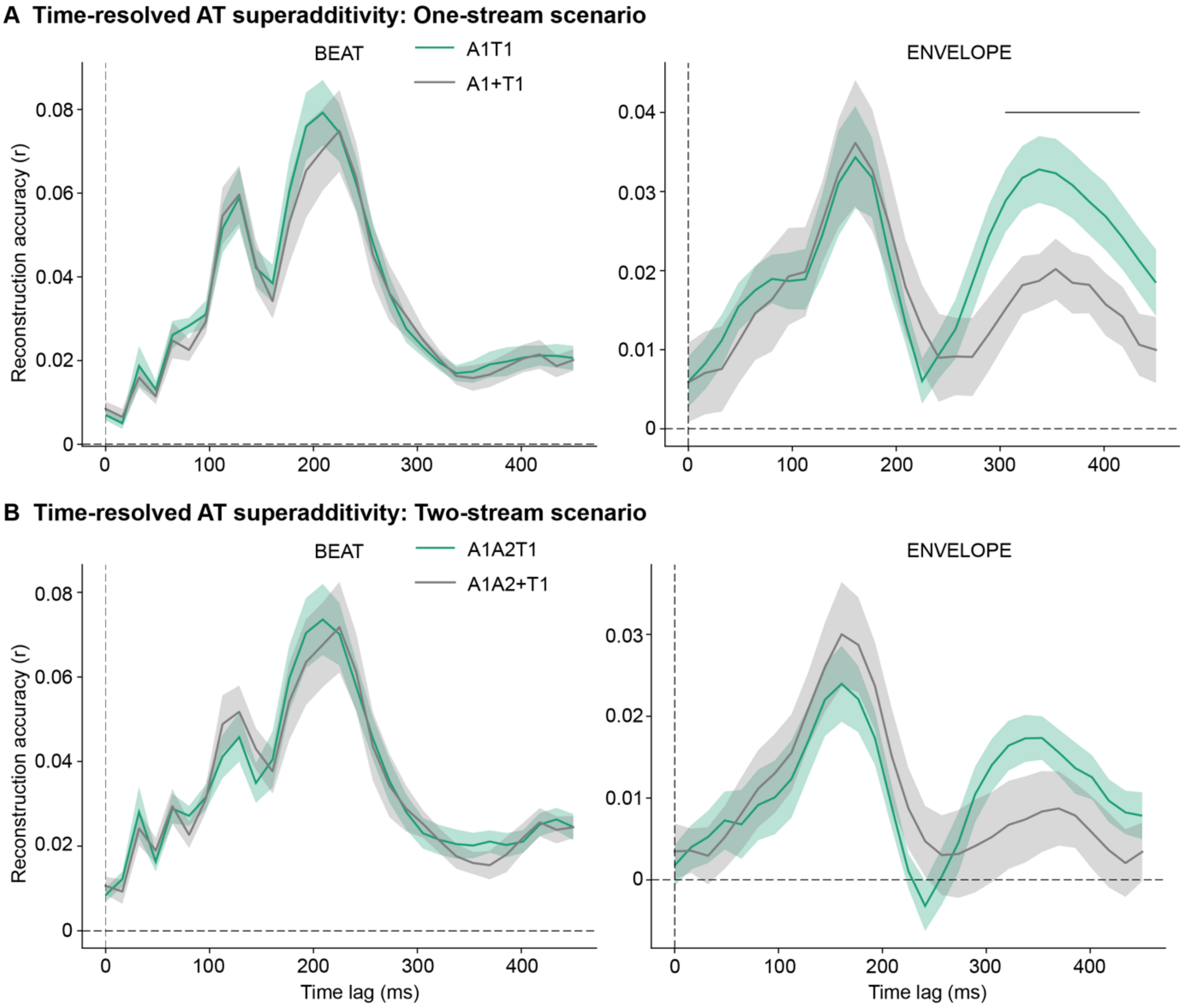


**Figure S2 | Time-resolved audiotactile superadditivity in one-stream and two-stream scenarios.** Across-participants' average time-resolved reconstruction of beat (left) and envelope (right) features. **A)** We plot results for audiotactile congruent (A1T1, green) against the sum of the respective unisensory auditory (A1) and tactile (T1) streams (A1+T1, grey) to visualize one-stream superadditive effects. **B)** We plot results for the audiotactile congruent stream (A1A2T1, green) against the sum of the respective unisensory auditory (A1A2) and tactile (T1) streams (A1A2+T1, grey) to visualize two-stream superadditive effects. Plots show mean (line) ± SEM (shaded area). For all plots, reconstruction accuracy is quantified in terms of Fisher-z-transformed Pearson's correlation (r) between the reconstructed and true feature time-series. Note that reconstruction accuracy is based on a decoding model trained on the unisensory auditory condition (A1) to examine selectively how auditory representations are influenced by concurrent tactile signals. The horizontal line shows the statistically significant temporal cluster for envelope ($p < 0.05$, corrected for multiple comparisons).

| | $d'$ | $bias$ |
|---|---|---|
| **One-stream scenario** | | |
| T1 | 1.26 (± 0.21) | 0.29 (± 0.09) |
| A1 | 1.25 (± 0.13) | 0.80 (± 0.08) |
| A1T1 | 1.69 (± 0.15) | 0.30 (± 0.09) |
| **Two-stream scenario** | | |
| T1 | 1.71 (± 0.21) | 0.04 (± 0.08) |
| A1A2 | 1.54 (± 0.12) | 0.79 (± 0.08) |
| A1*A2T1 | 1.91 (± 0.15) | 0.36 (± 0.07) |
| A1A2*T1 | 0.75 (± 0.09) | 0.94 (± 0.07) |

**Table S1 | Psychophysics results.** Across participants' mean (±SEM) detection sensitivity ($d'$) and decision strategy ($bias$) for one-stream and two-stream scenarios: one-stream auditory (A1); one-stream tactile (T1); one-stream audiotactile congruent (A1T1); two-stream auditory (A1A2); two-stream audiotactile with target in the congruent auditory and tactile streams (A1*A2T1) or in the incongruent auditory stream (A1A2*T1).
